# Interferon-α–Driven Stratification of B Cell Reveals Metabolic Reprogramming of Double Negative, Naive and Transitional cell subsets and Refines Molecular Classification in Sjögren’s Disease

**DOI:** 10.1101/2025.08.27.672530

**Authors:** Cristian Iperi, Akhiya Anilkumar Rekha, Gustav Arvidsson, Gunnel Nordmark, Eleonore Bettachioli, Marie Frutoso, Marta Alarcón-Riquelme, Jessica Nordlund, Ann-Christine Syvänen, PRECISESADS Flow Cytometry Study Group, PRECISESADS Clinical Consortium, Divi Cornec, Sophie Hillon, Christophe Jamin, Anne Bordron

**Affiliations:** LBAI, UMR1227, Univ Brest, Inserm, Brest, France; AltraBio SAS Lyon, France; Department of Medical Sciences, Molecular Precision Medicine, Uppsala University, Uppsala, Sweden; Science for Life Laboratory, Uppsala University, Uppsala, Sweden; Department of Medical Sciences, Rheumatology, Uppsala University, Uppsala, Sweden; CHU de Brest, Brest, France; GENYO, Centre for Genomics and Oncological Research Pfizer, University of Granada, Andalusian Regional Government, PTS Granada, Granada, Spain; Institute for Environmental Medicine, Karolinska Institutet, Stockholm, 171 69, Sweden

**Keywords:** B lymphocyte, transcriptomics, metabolism, Sjögren disease, Interferon signaling

## Abstract

Sjögren’s disease (SjD) is a chronic autoimmune condition marked by lymphocytic infiltration of exocrine glands and production of autoantibodies such as anti-SSA/Ro, anti-SSB/La, and rheumatoid factor. B lymphocytes play a central role in disease pathogenesis, driving autoantibody production and glandular damage, and contributing to lymphomagenesis. Despite promising therapies, no effective treatment is currently available, partly due to the biological and clinical heterogeneity of the disease. While interferon (IFN) signatures and B cell–related markers are used for patient stratification, their integration remains unexplored.

This study analyzed B cell transcriptional and metabolic profiles using bulk transcriptomic, clinical, and flow cytometry data from the PRECISESADS consortium, alongside public single-cell RNA-sequencing datasets. A B cell–specific IFN-α signature was established to stratify patients into IFN-positive and IFN-negative groups. Both showed reduced oxidative phosphorylation (OXPHOS) and translation in B cell subsets, suggesting a shared pre-metabolic state. IFN-positive patients, however, displayed additional features, including enhanced glycolysis, amino acid and lipid metabolism, autophagy, and NF-κB signaling. They also showed an expansion of IFN-activated naïve (Naive IFN), Transitional, and double-negative (DN) B cells, particularly DN2 and DN2_CXCR3 subsets, which have been linked in the literature to autoreactivity and lymphoma development.

The IFN signature in naïve B cells and DN2 correlated with elevated anti-SSA/Ro and anti-SSB/La titers, while only naïve B cells showed an association with increased histological focus scores. These findings support the relevance of a B cell–specific IFN signature in stratifying SjD patients and suggest new metabolic and transcriptional targets for disease monitoring and therapeutic development.

**Graphical abstract:** 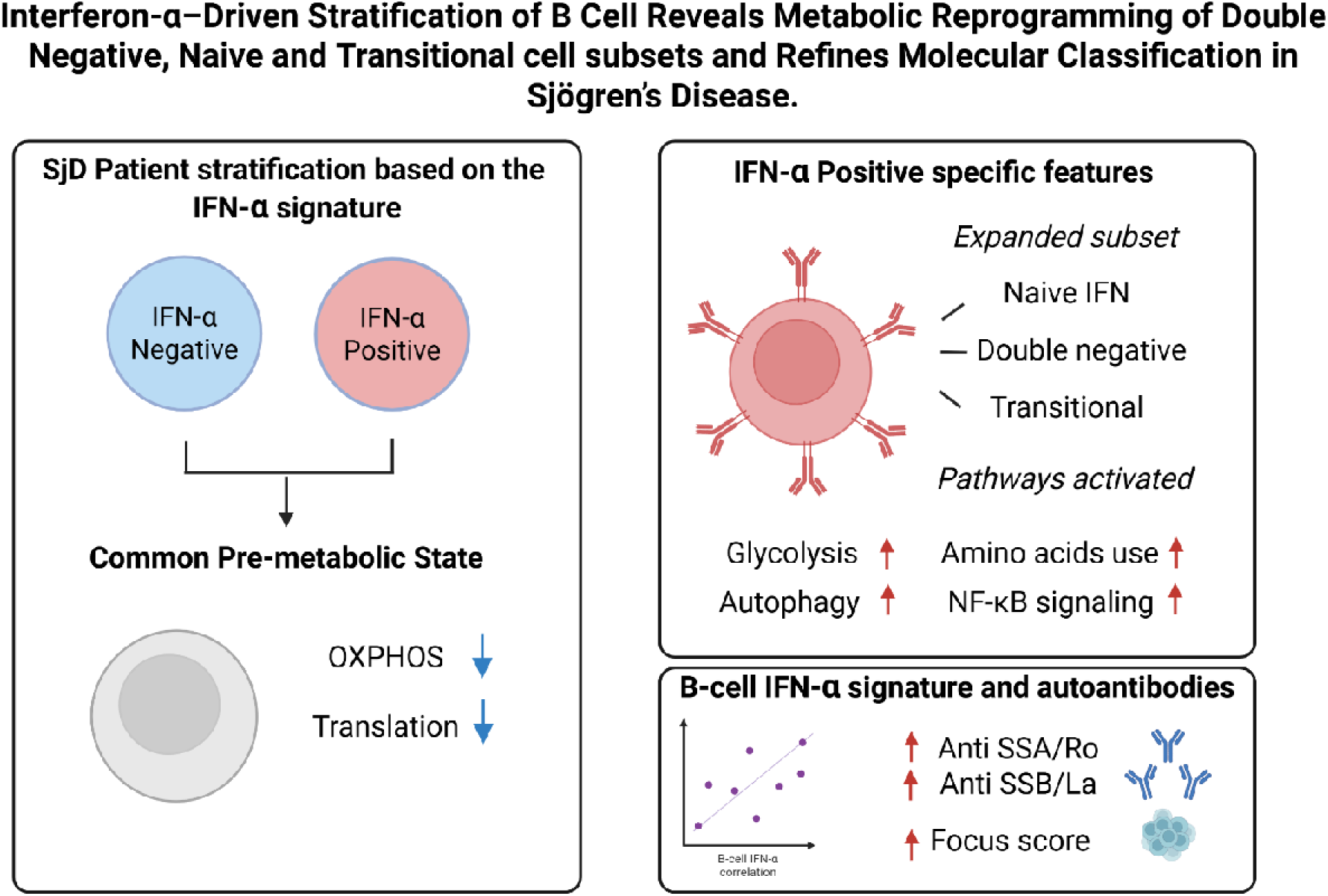

## 1. Introduction

Sjögren’s disease (SjD) is a chronic systemic autoimmune disease characterized by immune-mediated destruction of exocrine glands, particularly the salivary and lacrimal glands, leading to hallmark symptoms of dry mouth and dry eyes [1,2]. Serologically, the disease is strongly associated with circulating autoantibodies, including anti-Ro/SSA, which serves as a key diagnostic and prognostic marker, as well as anti-La/SSB and rheumatoid factor (RF) [2,3]. B cells are central to the pathogenesis of SjD, given their role in autoantibody production, their contribution to systemic manifestations, and their predominance among lymphocytic infiltrates in the salivary glands [4]. This infiltrate is associated with disease severity and progression and is commonly assessed using the focus score, a histological measure defined as the number of dense lymphocytic aggregates (foci of ≥50 lymphocytes) per 4 mm² of salivary gland tissue. [5]. Importantly, the presence of FcRL4^⁺^ B cells has been linked to the formation of lymphoepithelial lesions and the development of mucosa-associated lymphoid tissue (MALT) lymphomaC[6,7]. Although our understanding of SjD immunopathology has advanced, effective targeted treatments remain unavailable, and patients often continue to experience progressive symptoms and systemic complications [4]. This unmet clinical need is largely driven by the high heterogeneity of SjD, reflected in diverse autoantibody profiles (such as anti-Ro/SSA, anti-La/SSB, or seronegative cases), variable clinical manifestations (ranging from glandular dryness to systemic organ involvement), and differing levels of biological inflammation. This immunoclinical variability has significantly hindered the success of broad, common therapeutic strategiesC[8,9].

To address heterogeneity, patient stratification strategies have been increasingly explored, aiming to define more homogeneous subgroups for clinical research and treatment. Notably, stratification based on B lymphocyte characteristics has shown promise revealing that B cell–driven profiles are associated with more severe disease and may predict treatment responseC[8,10]. Parallel to this, another current stratification method involves measuring the type I interferon (IFN-α) transcriptional signature, an innate immune marker present in a substantial proportion of SjD patients [11–13]. The IFN signature is associated with heightened disease activity, elevated autoantibody titers, and increased risk of systemic complications [14]. Given the strong pathogenic role of both B cells and IFN signaling in SjD, integrating these two axes into a unified stratification framework could significantly enhance disease characterization and therapeutic targeting. B cells are both targets and mediators of IFN signaling [15], suggesting that a B cell–specific IFN signature might more accurately reflect the active autoimmune processes in individual SjD patients.

Support for this integrated approach comes from studies in systemic lupus erythematosus (SLE), an autoimmune disease that is immunologically and biologically similar to SjD [16]. In SLE, IFN-α is known to drive B cell activation, promote the loss of tolerance, and support the expansion of autoreactive B cell subsets C[17,18]. Notable IFN-driven changes in B cell populations include enhanced pro-inflammatory and survival features in transitional B cells, direct activation and differentiation of naïve B cells, and presence of the double-negative (DN) B cell subset [18–20].

Given the need to better capture the clinical and biological heterogeneity of SjD through more targeted stratification strategies, particularly those involving B cell, this study aimed to comprehensively dissect the diversity of B cell states and their relationship to interferon exploring the transcriptional and phenotypic profiles of B cells in depth. To this end, bulk transcriptomic, clinical, and flow cytometry data from the PRECISESADS consortium [16], along with publicly available single-cell RNA-sequencing datasets [21], were used. This integrative approach aimed to unveil the molecular underpinnings of B cell–driven heterogeneity in SjD and to evaluate the potential of a B cell–specific IFN signature as a clinically meaningful tool for patient stratification.

## 2. Material and methods

### 2.1 Samples and cohort selection

The cross-sectional dataset from the European PRECISESADS cohort (number NCT02890121 in https://ClinicalTrials.gov) aimed to reclassify the autoimmune diseases by the molecular signature including clinical and multi-omics information. The patient selection and quality control have previously been described in detail [16]. PRECISESADS adhered to the standards set by International Conference on Harmonization and Good Clinical Practice (ICH-GCP), and to the ethical principles written in the Declaration of Helsinki (2013). Each patient signed an informed consent prior to study inclusion. The Ethical Review Boards of the 19 participating institutions approved the protocol of the cross-sectional study. This study was a pre-planned substudy to be specifically conducted in the SjD population, defined by the 2002 American-European Consensus Group classification criteria [22].

### 2.2 Transcriptome data generation

#### Bulk dataset

Briefly, peripheral blood from 41 patients and 27 healthy controls (CTRLs) from PRECISESADS cohort were collected. B cells were then isolated using REAlease CD19 Microbead Kit (Miltenyi Biotec), the purity verified by flow cytometry and the libraries obtained by TruSeq Stranded mRNA HT kit (Illumina). The samples were then sequenced using a NextSeq 500 with an average coverage per sample of 29.25 Million reads. The FastQ files were aligned to the UCSC *Homo sapiens* reference genome (hg19) and annotated to GENCODE 19 with STAR v2.5.2 [5] using two-pass mapping strategy with default parameters. Gene quantification was performed using RSEM v1.2.31 [6].

#### Single cell dataset

The single cell dataset was downloaded from the GEO database (GSE214974). It contained the B-cell transcriptomes from four CTRLs and 24 patients with SjD, all females. One sample (SJD-024) was excluded because of low B lymphocytes according to the original manuscript [21]. The samples SJD-001 and SJD-014 were removed due to insufficient quality (**Table S1**).

### 2.3 Analysis of bulk B-cell purity and metadata

B-cells were isolated by CD19 positive selection (Miltenyi) and the purity checked by flow cytometry. After extraction, the B-cell RNA purity of SjD and CTRL samples was evaluated using Microenvironment Cell Populations-counter [23] on R with default parameters, a tool based on immune cell-specific gene signatures. Samples with digital purity lower than 90% were excluded from the subsequent analysis. The final number of samples included 28 SjD and 23 CTRL samples **(Table S2).** The associated clinical metadata were statistically compared by anova test and t-test, and the general characteristics reported in **Table 1**. The cytokines, autoantibodies and flow cytometry data generation have been extensively described in detail in the PRECISESADS publication [16]. For the single-cell data, patient-associated metadata are available in the Table 1 of the original publication describing the dataset [21].

**Table 1.** Healthy controls (CTRL) and Sjögren disease (SjD) patients characteristics in PRECISESADS data.

|  |  | CTRL<br>(n=23) | SjD<br>(n=28) |
| --- | --- | --- | --- |
| Age | Mean ± SD | 49.35 ± 10.98 | 53.82 ± 13.56 |
| ENA (pos≥1) | n (% pos) | 0 (0%) | 14/15 (93.3%) |
| Anti-SSA (pos≥10) | n (% pos) | 0 (0%) | 14/15 (93.3%) |
| Anti-SSA-52 (pos≥10) | n (% pos) | 0 (0%) | 6/7 (85.7%) |
| Anti-SSA-60 (pos≥10) | n (% pos) | 0 (0%) | 5/7 (71.4%) |
| Anti-SSB (pos≥10) | n (% pos) | 0 (0%) | 5/15 (33,3 %) |
| Anti-MPO (pos≥40) | n (% pos) | 0 (0%) | 0 (0 %) |
| Anti-PR3 (pos≥40) | n (% pos) | 0 (0%) | 0 (0 %) |
| Anti-Centromer (pos>10) | n (% pos) | 0 (0%) | 2/13 (15,38 %) |
| C3 (g/L) | Mean ± SD | 0.91 ± 0.42 | 1.14 ± 0.41 |
| C4 (g/L) | Mean ± SD | 0.16 ± 0.08 | 0.16 ± 0.07 |
| RF (UI/mL) | n (% pos) | 1 (4,3 %) | 5 (33,3 %) |
| Lambda chain (mg/L) | Mean ± SD | 6.75 ± 5.93 | 15.69 ± 8.08 |
| Kappa chain (mg/L) | Mean ± SD | 7.35 ± 5.00 | 21.23 ± 19.97 |
| Anti-DNA and nucleosome (UI/mL) | Mean ± SD | 15.87 ± 14.15 | 59.49 ± 113.47 |
| Cigarette smoking | n (%) | 3 (13.0%) | 3 (10.7%) |
| Sicca syndrome | n (%) | 0 (0%) | 25 (89.3%) |
| Raynaud's phenomenon | n (%) | 0 (0%) | 7 (25.0 %) |
| Hypertension | n (%) | 0 (0%) | 10 (35.7%) |
| History of cancer | n (%) | 0 (0%) | 2 (7.1%) |
| Use of steroids | n (%) | 0 (0%) | 6 (21.4%) |
| Use of immunosuppressant | n (%) | 0 (0%) | 4 (14.2%) |
| Hydroxychloroquine | n (%) | 0 (0%) | 14 (50.0%) |
| Involvement central nervous system | n (%) | 0 (0%) | Present/Past = 3 (2.0%) |

n: Number of patients. If the data are missing from a variable the number of patients with the available information are indicated as denominator; pos: positivity.

### 2.4 Flow cytometry data analysis

The PBMCs flow cytometry data harmonised according to previous works were extracted from the matched B cell samples to assess B cell subpopulation changes. In brief the gating strategy included a panel including IgD (FITC), CD267 (PE), CD5 (PC5.5), CD19 (APC), CD24 (APC-AF-750), CD38 (PB). More information is available from the original works [13,24]. The percentage of each B cell subset within total B cells was compared between IFN-pos, IFN-neg, and CTRL groups using the Wilcoxon test.

### 2.5 Single-cell data clustering and analysis

The analysis of the single cell was replicated from the original manuscript [21]. Seurat R package v4.3.0 was used to analyze the 255 462 unique cellular barcodes removing cells with ≥ 20% mitochondrial UMIs (686 cells), ≥ 1 % hemoglobin UMIs (4365 cells) or ≤ 5% ribosomal UMIs (5826 cells), as the original manuscript. The cells with the number of detected features or total UMI counts belonging to the 98th percentile in each sample were discarded (15330 cells). The counts were normalized by LogNormalization() function and the most variable features selected (n = 4000) and used to compute the 50 first principal components by RunPCA(). The batch correction per sample was made by RunHarmony() and then the RunUMAP() was run using the harmony dimensionality reduction with the following parameters: dims = 1:50, metric = “correlation”, min.dist = 0.4, spread = 0.5, n.neighbors = 30, repulsion.strength = 0.4, negative.sample.rate = 50 and n.epochs = 100. FindAllMarkers() extracted the top featured with the parameter settings: min.pct = 0.1, min.cells.feature = 0.1, max.cells.per.ident = 50, only.pos = TRUE and logfc.threshold = 0. Twenty-four clusters were detected, including some containing T lymphocytes and monocytes. These clusters were excluded for further analysis. The same features selection, PCA, batch correction and UMAP were re-runned on the filtered B cells and the markers identified by FindAllMarkers() with the parameter settings: min.pct = 0.1, min.cells.feature = 0.1, max.cells.per.ident = 200, only.pos = TRUE and logfc.threshold = 0.1.

### 2.6 IFN-**α** classification in bulk and single cell B-cell samples

Patients with SjD were clustered using an IFN-α score established based on the Kirou score [25] approach. This approach measures the IFN-α response using the transcriptome of 20 genes [26] (*SIGLEC1*, *IFIT3*, *IFI6*, *LY6E*, *MX1*, *USP18*, *OAS3*, *IFI44L*, *OAS2*, *IFIT1*, *EPSTI1*, *ISG15*, *RSAD2*, *HERC5*, *OAS1*, *IFI44*, *SPATS2L*, *PLSCR1*, *IFI27*, and *RTP4*) in addition to six common IFN-α-γ-associated genes (*EIF2AK2*, *GBP1*, *IRF1*, *SERPING1*, *CXCL10*, *FCGR1A*) [27]. Gene expression levels from bulk RNA seq, were quantified in terms of raw reads normalised to the total number per sample, and the score was calculated as the expression of each gene (g) for each SjD sample (s) with the mean expression divided by the standard deviation of the CTRLs as follows:

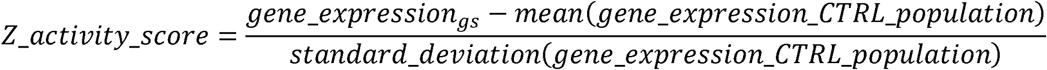

The higher the Z interferon activity score (Z-score) for a particular gene compared to the CTRLs, the higher the IFN-α activation of that gene. By IFN gene Z-scores, samples were grouped using the hierarchical clustering of the Complexheatmap package v2.12.0 [28] on R by Euclidean distance and ward.D2 clustering method. This analysis stratified B cells from SjD patients into four distinct clusters: three IFN-α–positive clusters (clusters 1, 3, and 4) and one IFN-α–negative cluster (cluster 2), resulting in 18 IFN-α–positive (IFN-pos) and 10 IFN-α–negative (IFN-neg) samples. Similarly, hierarchical clustering was applied to pseudobulk single-cell RNA-seq data prior to Z-score activity calculation, mirroring the approach used for bulk B cells. Four clusters were identified: three with high IFN-α expression (clusters 2, 3, and 4) and one with low-to-no expression (cluster 1). To maintain consistency with the bulk analysis, only the IFN-α–negative subset of cluster 1 was included in the IFN neg group, while the remaining samples were assigned to the IFN pos group. This resulted in 17 IFN pos and 4 IFN neg samples. The Chaussabel modules M1.2, M3.4, and M5.12 were calculated in the original PRECISESADS study on SjD using whole blood transcriptome data, based on the expression of genes such as PRKR, IFIT1, and IFI44. [13].

### 2.7 B lymphocytes subpopulation annotation in single cell

The annotation was performed to retrieve the one from the original paper. ScPred (v1.9.2) [29] annotated the single-cell data by a model trained on the sorted B cells provided in the study of Stewart et al [30]. This allowed the identification of two populations of CD19^+^CD27^+^IgD^-^classical memory (C-mem), CD19^+^IgD^+^CD27^+^ IgM memory (M-mem), four populations of CD19^+^IgD^-^CD27^-^Double Negative (DN), and a single population of CD19^+^IgD^+^CD27^-^CD10^+^ Transitional and CD19^+^IgD^+^CD27^-^CD10^-^Naïve B cells. The clusters were then manually annotated by comparing the markers detected in the original paper and the ones calculated by FindAllMarkers(). The B cell subpopulation for the IFN pos, IFN neg and CTRLs were reported in **Table S3**.

### 2.8 Differential gene expression analysis

All differential gene expression (DGE) analyses provided raw gene counts as input for DESeq2 [31], used with default parameters. In the case of single cell data, the pseudo-bulk matrices were calculated for each B cell subtype within each patient and used as row gene counts. In bulk analyses, sex, age, and the most common treatments, steroids and hydroxychloroquine, were included in the model as covariates to account for potential confounding effects. The threshold values to identify the DEGs were set to |Log2FC| > 0.5 and false discovery rate (FDR) <0.05. Nominal *p*-values were adjusted for multiple testing using the FDR method [32].

### 2.9 Gene set enrichment analysis

The gene set enrichment analysis was performed using GSEA [33] by the fgsea R package V1.34.0. The matrix counts were uploaded in R and the Biological Processes (BP) V.2024.1 library was used. The EnrichR [34] tool v2022 was also used, uploading the genes identified as upregulated and downregulated in the DGE analysis separately. For both, only processes with a Benjamini–Hochberg adjusted p-value < 0.05 were considered.

### 2.10 Trajectory analysis by Slingshot

The single cell trajectory analysis was performed on Slingshot [35] V2.14 on a subset of 10.000 cells with default parameters. Five trajectories were identified. The first and the last trajectories were analysed, respectively the one connecting transitional to DN2 and the one within DN population. By Tradeseq [36] v1.20 with default parameters the differentially expressed genes were identified and the relevant biological pathway identified by EnrichR. As only Trajectory 1 showed enrichment in biologically meaningful pathways, a heatmap was generated displaying genes associated with the selected pathways identified by the associationTest() results. These pathways included IFN signaling (R-HSA-913531), cellular stress (R-HSA-2262752), B cell activation (GO:0050853, GO:0042113, GO:0030888), and metabolic processes such as glycolysis (R-HSA-70326), amino acid activation (R-HSA-71291), and lipid metabolism (R-HSA-556833).

### 2.11 Correlation analysis

The Kirou IFN score, derived from whole blood transcriptomic data as reported by Soret et al. [13], was compared with the B cell–specific IFN score obtained from bulk B cell transcriptomes. Both scores were assessed for their correlation with SjD patient autoantibody levels (measured by ELISA). Pearson correlation analyses were conducted to evaluate similarities and differences in their associations with serological markers. To refine the analysis, subset-specific IFN correlations were performed using single-cell transcriptomic data. For each B cell subset, the mean expression of IFN-α–related genes was calculated and correlated with autoantibody levels as measured by Arvidsson [21] et al. by an addressable laser bead immunoassay (Luminex) and nephelometry, respectively, and clinical parameters.

## 3. Results

### 3.1 B Cell IFN-α Stratification Reveals Distinct Transcriptional Profiles in SjD Patients

To initiate the characterization of SjD patients based on the B cell IFN-α signature, transcriptomic profiling of isolated B cells from the PRECISESADS cohort was performed. The B cell IFN-α signature was assessed using a validated panel of 26 interferon-stimulated genes (ISGs), previously applied to SLE patients based on B cell IFN-α activity [26,27]. For each gene, Z-scores were calculated using the mean and standard deviation of expression in the CTRL population as a reference, and used for hierarchical clustering. This analysis stratified SjD patient B cells into four distinct clusters: three IFN-α– positive groups (clusters 1, 3, and 4) and one IFN-α–negative group (cluster 2)(**Figure 1A; Figure S1**). Despite variations in IFN-α intensity, IFN-α–positive clusters were merged to increase cohort size and enhance their robustness and subsequent analyses. This stratification was further supported by an independent IFN-α scoring approach, calculated from PBMC transcriptomic data in the PRECISESADS cohort, as described previously [13]. IFN-α module scores, derived from Chaussabel’s modular framework, included module M1.2 (specific for IFN-α), M3.4 (responsive to both IFN-α and IFN-γ), and M5.12 (specific for IFN-γ) (**Figure 1B**).

**Figure 1:**
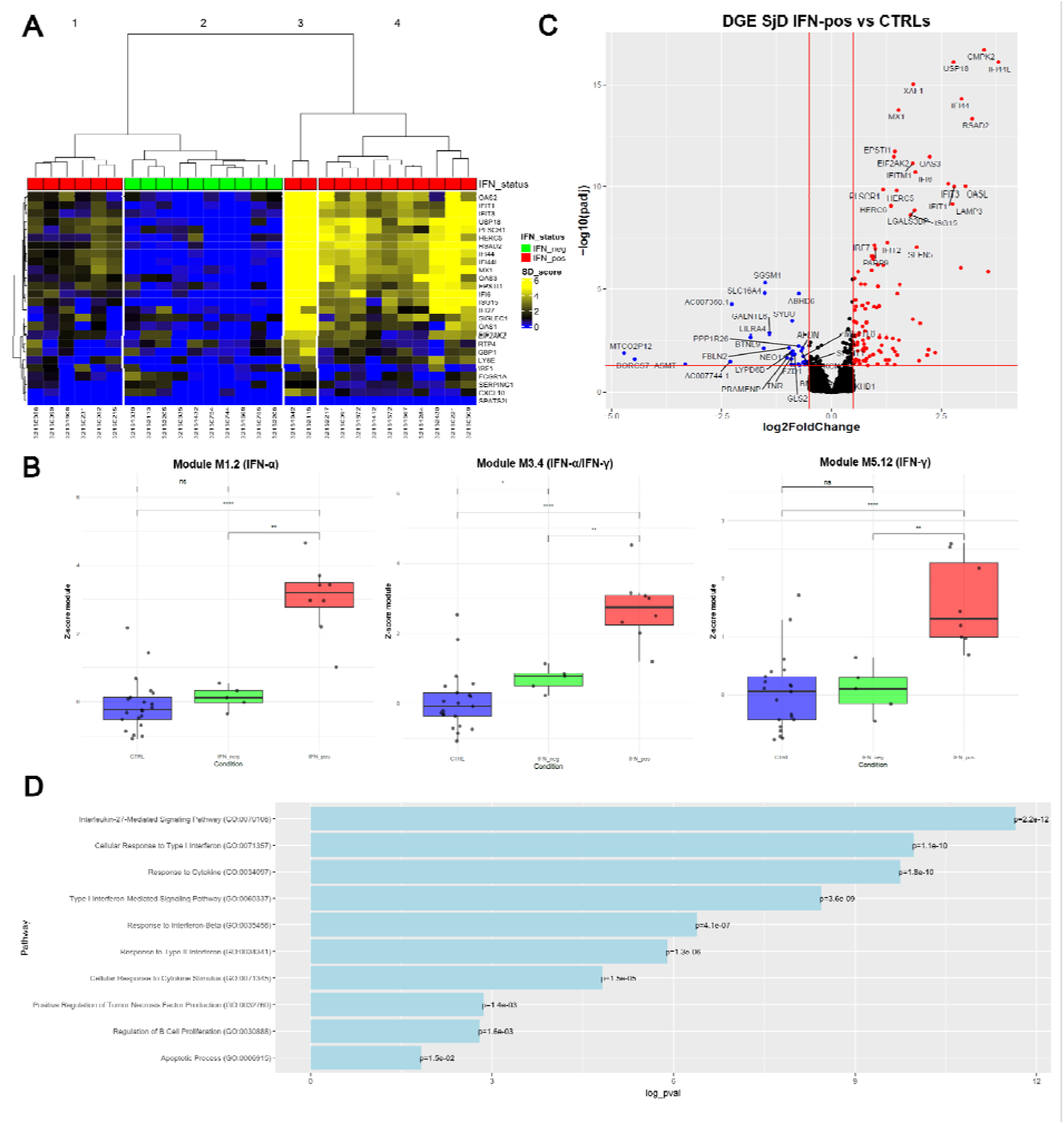
IFN-α–Based Stratification Reveals Distinct Transcriptomic and Functional Pathway Signatures in SjD B Cells. **A)** Hierarchical clustering of SjD patients based on the B cell IFN-α signature from the PRECISESADS dataset. The heatmap displays IFN status: IFN–pos patients are shown in red, and IFN-neg patients in green. Columns represent individual samples; rows correspond to IFN-α signature genes. Each cell indicates the gene expression Z-score relative to the CTRL group. **B)** Scatter plot showing Z-scores of Chaussabel’s IFN-related modules (M1.2: IFN-α–specific, M3.4: IFN-α/γ–responsive, M5.12: IFN-γ– specific) in matched PBMC samples, used to validate the IFN signature observed in B cells. **C)** Volcano plot of the differential gene expression analysis comparing IFN-pos SjD patients to CTRLs in the PRECISESADS bulk B cell dataset. Significantly upregulated genes are shown in red, and downregulated genes in blue. The top 25 genes in each direction are labeled. Red lines indicate significance thresholds: |log₂FC| > 0.5 and false discovery rate (FDR) < 0.05. **D)** Chart plot of significantly enriched pathways identified by EnrichR analysis from SjD IFN-pos compared to CTRLs. The X-axis represents the –log₁₀ of the adjusted p-value, while the Y-axis lists the pathways ranked by statistical significance.

To investigate differences in the B cell transcriptomic profiles of IFN-stratified SjD patients, a DGE analysis on the interferon-α positive (IFN-pos) (**Figure 1C**) and interferon-α negative (IFN-neg) patients (**Figure S2**) was performed using the CTRLs as reference. A total of 104 genes were found to have a higher expression and 33 a lower expression in the IFN-pos group compared to CTRLs, while no significant differences were observed between the IFN-neg group and CTRLs. Among the higher expressed genes, ISGs such as IFI44L, RSAD2, and OAS3 were identified, as well as genes associated with B cell activation, including TNFSF13B (BAFF) and CD38. Enrichment analysis using EnrichR (GO:BP 2025, Reactome 2024) confirmed the involvement of pathways related to type I interferon signaling (GO:0034340), response to virus (GO:0051607), apoptotic processes (GO:0006915), positive regulation of B cell proliferation (GO:0030890, GO:0050671), and IL-27 and IL-6 signaling pathways (GO:0070106, R-HSA-1059683) (**Figure 1D**). The 33 lower expressed genes did not show enrichment in any significantly associated biological pathways. No significant differences in gene expression were observed between the IFN-neg group and CTRL (**Figure S2**). To further investigate pathway-level alterations, GSEA was performed by the GSEA V4.4.0 program on the gene expression rankings obtained by the IFN-pos and IFN-neg DGE analysis. The analysis confirmed that the IFN-pos group was enriched for pathways related to type I interferon signaling, antiviral responses, B cell-mediated immunity, and apoptosis. In contrast, the IFN-neg group did not show enrichment for any gene sets. The GSEA, DGE and EnrichR results are reported in **Supplementary Material 1**.

### 3.2 Flow Cytometry Validates IFN-**α**–Based B Cell Stratification in SjD

As the observed IFN signatures were derived from bulk B cell transcriptomic data, it was necessary to assess whether these transcriptional profiles were associated with specific B cell subsets. To this end, flow cytometry data generated from PBMCs were analyzed to define B cell populations. A gating strategy was applied to isolate CD19^⁺^ B cells (**Figure 2A**), and the frequencies of defined subsets were compared across IFN-stratified patient groups. A reduction in memory B cell populations was observed in both IFN-positive and IFN-negative SjD patient groups compared to controls. This significant reduction encompassed multiple subsets, including activated memory B cells, such as CD19^⁺^IgD^⁻^CD27^⁺^ class-switched memory cells and CD19^⁺^IgD^⁺^CD27^⁺^unswitched memory cells, as well as resting memory subsets, including CD19^⁺^CD24^⁺^CD38^-^resting memory B cells, CD19^⁺^IgD^⁺^CD38^⁻^ Bm5 cells, and CD19^⁺^IgD^⁺^CD38^⁻^ early Bm5 cells.

**Figure 2:**
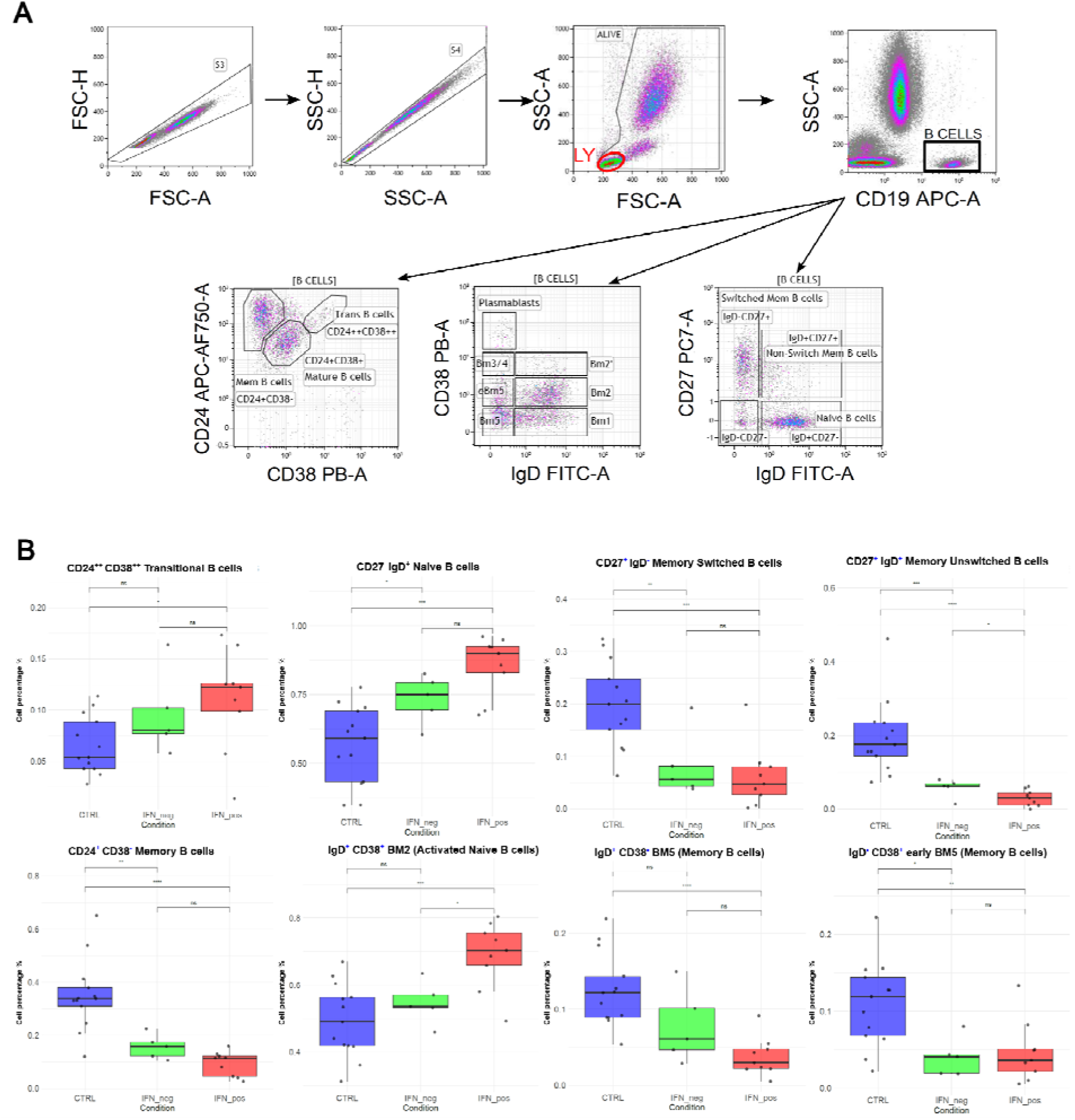
IFN-α–Based Stratification Reveals Distinct B cell subpopulations in SjD patients. **A)** Gating removed cell doublets and identified the B cell population. The B cell subpopulations were subsequently defined using combinations of IgD, CD27, CD24, and CD38 markers. **B)** Boxplots of flow cytometry data across IFN-α groups. The X-axis indicates IFN-α groups (IFN-pos, IFN-neg, and CTRL), and the Y-axis shows B cell subsets percentage within total B cells. Sample size: *n* = 27 (9 IFN-positive, 5 IFN-negative, 13 CTRL).

Among the distinct features observed across IFN-stratified groups, the IFN-pos group exhibited a marked expansion of CD19^⁺^CD24^⁺^CD38^⁺^ transitional B cells and CD19^⁺^IgD^⁺^CD27^⁻^ naïve B cells, with a less pronounced increase in the IFN-neg group. The transitional expansion was supported by the previously observed CD38 in bulk transcriptomics data. Furthermore, the IFN-pos group displayed a distinctive expansion of CD19^⁺^IgD^⁺^CD38^⁻^activated naïve B cells subset, consistent with an expansion of naïve B cells. The gating strategy used to define B cell subpopulations is shown (**Figure 2A**), along with the corresponding percentages of each B cell subset (**Figure 2B**).

### 3.3 IFN-**α**–Driven B Cell Subpopulation Changes Confirmed by Single-Cell Analysis

In the previous sections, it was shown that SjD patients display varying intensities of a B cell–intrinsic IFN-α transcriptional signature. Stratifying donors based on this signature revealed differences in the frequencies of specific B cell subsets, particularly unswitched memory cells and activated naïve cells. Both subsets showed significant changes between IFN-positive and IFN-negative groups. This raised key questions: do specific biological changes accompany stronger IFN-α signatures, and could these changes help explain disease heterogeneity? From the literature, SSA and SSB status were positively associated with the IFN-α signature, particularly in monocytes, while BAFF was associated with disease severity [21,37,38]. However, much less is known about how the IFN-α signature manifests in B cells. Bulk transcriptomic data provide an overview of transcriptional differences but lacks the resolution to capture shifts in B cell subset abundance or phenotype. To overcome this limitation, a publicly available single-cell RNA-sequencing dataset of peripheral B cells from 24 SjD patients and 4 controls (G. Arvidsson et al. [21]]) was analyzed. This high-resolution dataset enabled a deeper exploration of B cell heterogeneity and the cellular dynamics linked to IFN-α signatures, offering insights into potential mechanisms driving disease progression.

To determine whether the stratification observed in bulk data could be recapitulated, pseudobulk RNA-seq data from all B cells were hierarchically clustered using the same approach as in bulk analysis. This confirmed the reproducibility of the IFN-based stratification identifying both IFN-pos and IFN-neg patient clusters (**Figure 3A; Figure S3**). The Cluster 1 contained two subclusters: one composed of strictly IFN-α negative patients, and another including patients with low, but detectable, IFN-α activity. Despite their lower expression levels, the latter were grouped within the IFN-pos category due to their overall transcriptional profile. As for bulk data, the different IFN-α intensities of the clusters having detectable IFN-α were included into the IFN-pos to increase the sample size and statistical power. Furthermore, the IFN-α gene expression patterns mirrored those observed in bulk RNA-seq.

**Figure 3:**
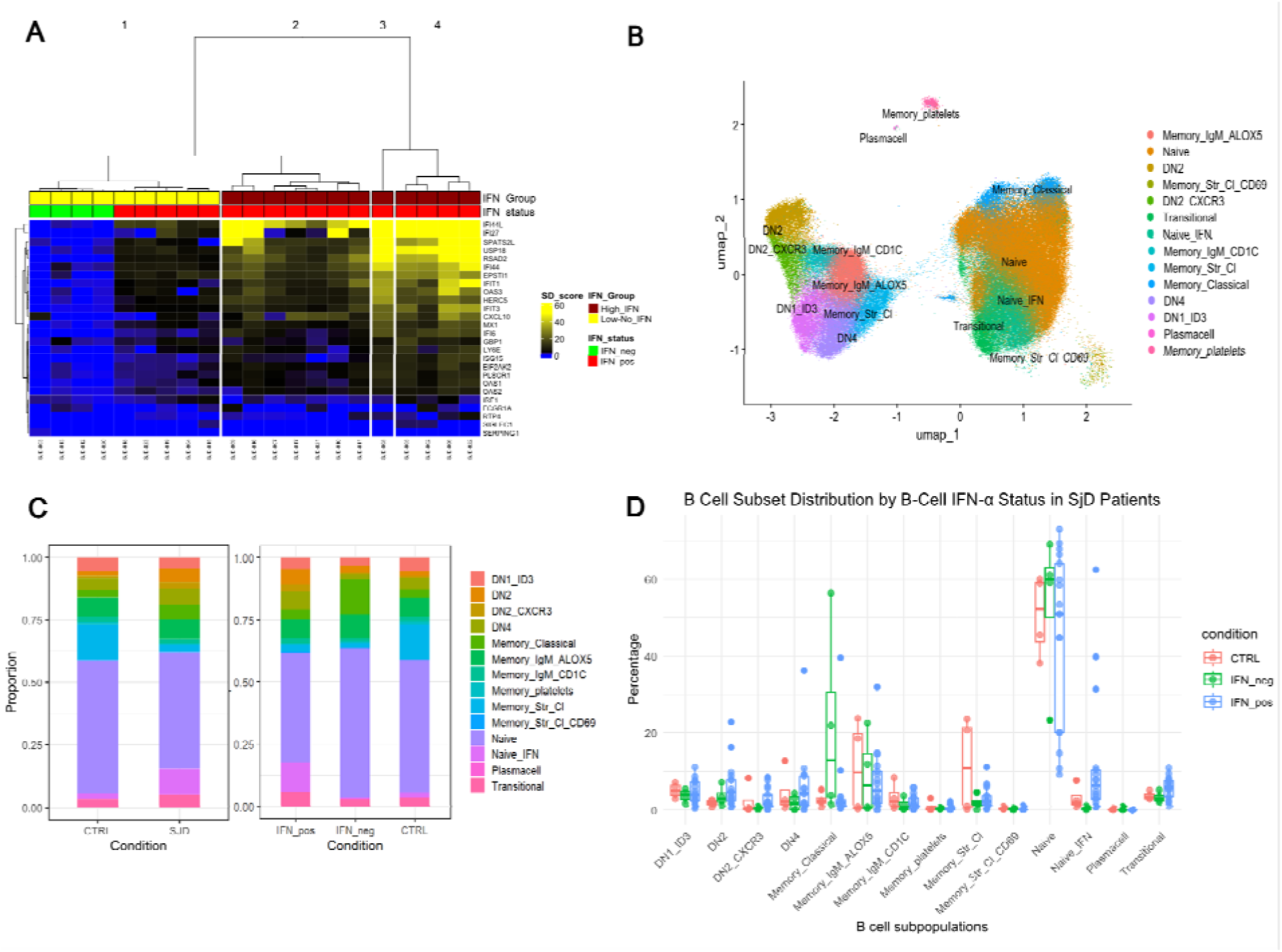
Single-cell analysis reveals IFN-α–driven heterogeneity and B cell subpopulation shifts in SjD patients. **A)** Clustering of public single cell data by hierarchical clustering. The heatmap displays IFN-α groups corresponding to the identified clusters: High IFN patients are shown in dark red, and Low/No IFN-α patients in yellow. A second annotation indicates IFN status, with IFN-α–positive patients in red and IFN-α– negative patients in green. Columns represent individual samples; rows represent IFN-α signature genes. Each cell shows the gene expression Z-score relative to the CTRL group. **B)** UMAP plot of the public single-cell dataset, with B cell subpopulations highlighted in different colors. **C)** Bar plot showing the proportions of B cell subpopulations in the public single-cell data. The left panel compares CTRL and SjD groups, while the right panel compares CTRL with the IFN-α–stratified groups. (X-axis: CTRL and IFN-α groups; Y-axis: proportion of each subpopulation) **D)** Boxplots of B cell subpopulation within IFN-α groups. (X-axis: B cell subpopulation; Y-axis: cell percentage). The CTRL group is shown in red and the IFN-neg and IFN-pos group respectively in green and blue.

To explore B cell heterogeneity in greater detail, single-cell data were clustered using the same method described in the original manuscript. A total of 24 clusters were annotated based on the original classification, including transitional, plasmacell, memory (Memory_Classical, Memory_IgM_ALOX5, Memory_IgM_CD1C, Memory_platelets, Memory_Str_Cl, Memory_Str_Cl_CD69), naive (Naive, Naive_IFN), and double-negative subsets (DN1, DN2, DN2_CXCR3, DN4). The analysis then focused on B cell subpopulations (**Figure 3B; Table S2)**, comparing their relative abundance and expression profiles across IFN-α–defined groups (**Figure 3C; Table S2)**.

The Naive IFN subpopulation, including Naive expressing ISG genes, was expanded in the IFN-pos group, consistent with the overall naive B cell and activated B cell expansion observed previously in the PRECISESADS flow cytometry data. A gradual increase in the DN2 population was observed across the IFN-neg to IFN-pos groups, accompanied by a concomitant expansion of the DN2 CXCR3^⁺^ and DN4 subpopulation. In agreement with the PRECISESADS flow cytometry data, classical memory B cells including unswitched memory cells (Memory_IgM_ALOX5, Memory_IgM_CD1C) were progressively reduced from IFN-neg to IFN-pos patients compared to CTRLs. Conversely, transitional B cells were increased in the IFN-pos group (**Figure 3C, 3D**).

### 3.4 Metabolic Reprogramming of B Cells Revealed by Single-Cell Analysis

To further dissect the heterogeneity of SjD, the transcriptional landscape of B cell subpopulations was examined after stratifying patients by B cell IFN-α signature status. Specifically, patients with IFN-pos and IFN-neg profiles were compared to CTRLs. The number of differentially expressed genes across all B cell subsets was higher in the IFN-pos group compared to the IFN-neg group. As expected, a strong type I IFN signature was consistently detected in all B cell subpopulations of the IFN-pos group, accompanied by upregulation of autophagy-related pathways. Notably, NF-κB signaling and apoptotic processes were enriched in classical memory B cells, DN1_ID3, and DN4 subsets within the IFN-pos group. Also, glycosylation pathways were significantly upregulated in IFN-pos classical memory, DN2, DN2_CXCR3, and DN4 subsets, with mannosylation processes exclusively enriched in DN2 and transitional B cells (**Figure 4A; Figure S4**). In contrast, the IFN-neg group demonstrated a marked reduction in immunoglobulin-related processes in DN2_CXCR3, memory IgM, stressed classical memory, transitional, and naïve IFN B cells (**Figure 4B; Figure S4**). Subset-specific features were observed in DN1_ID3 cells, with increased purinergic nucleotide signaling in both IFN-α groups, and increased sphingolipid metabolism exclusively in the IFN-pos group. Furthermore, DN2_CXCR3 cells exhibited specific expression of FcRL4 in 3.4% of the subpopulation, a feature linked to pro-lymphoma B cell populations (**Figure S5**).

**Figure 4:**
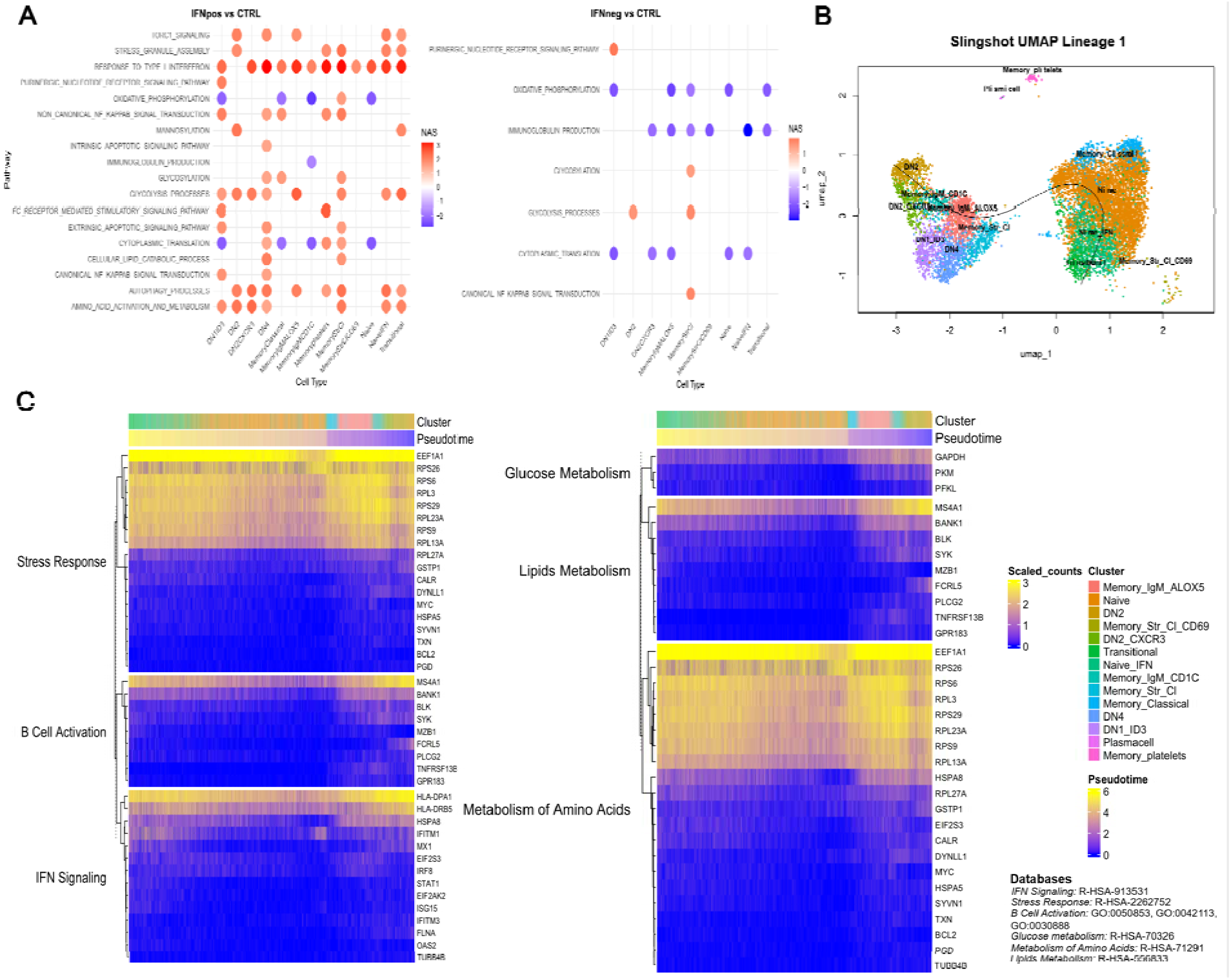
Interferon-α–Associated Pathways and Trajectories Reveal B Cell Activation and Metabolic Remodeling. **A)** Dot plot of enriched GSEA pathways in B cell subtypes comparing IFN-α–positive and IFN-α–negative patients versus CTRLs. Pathways are shown on the y-axis and B cell subtypes on the x-axis. Red indicates upregulated pathways; blue indicates downregulated pathways. Dot size reflects the normalized enrichment score (NES). If a B cell subpopulation is missing no significant processes were identified. **B)** Trajectory analysis of B cell maturation using Slingshot on Lineage 1, visualized on a UMAP plot with inferred developmental progression. **C)** Heatmap of B cell subtypes ordered by pseudotime along Trajectory 1. The x-axis shows genes identified as trajectory-relevant by associationTest() and involved in selected Reactome and Gene Ontology pathways: stress response (R-HSA-2262752), B cell activation (GO:0050853), IFN signaling (R-HSA-913531), glucose metabolism (R-HSA-70326), lipid metabolism (R-HSA-556833), and amino acid metabolism (R-HSA-71291). The expression of these trajectory-relevant genes was log1p-scaled, and the values were represented on a blue-to-yellow scale, from high to low, respectively. The same color scheme was applied to the pseudotime visualization.

Metabolic reprogramming represented a prominent feature across both IFN groups. A consistent B cell subpopulation downregulation of oxidative phosphorylation (OXPHOS) and translational processes was observed, distinguishing SjD patients from CTRLs regardless of IFN-α status. Glycolysis and amino acid metabolism were broadly activated in the IFN-pos group across all DN, Naive IFN and transitional subpopulations while in the IFN-neg group, glycolysis showed specific enrichment only in DN2 and stressed classical memory B cells. Lipid metabolism was also enhanced in the IFN-pos group, with increased lipid-related pathways in DN1_ID3 cells and lipid oxidation in naïve IFN, DN4, classical memory, and stressed classical memory B cells. To support this elevated metabolic activity within the IFN-pos group, mTORC, a master regulator of anabolic metabolism, was observed activated in transitional, naïve, memory IgM, DN2, and DN4 B cells (**Figure 4A**).

### 3.5 IFN-α sustain the Maturation toward Double-Negative B Cell Subpopulations

Stratification of SjD patients by IFN-α status in B cells revealed significant alterations in B cell subpopulation frequency, gene expression, and metabolic activity. The DN compartment emerged as a key site of IFN-α–driven changes, characterized by both expansion in the IFN-pos group and a distinct metabolic signature, indicating a potential role in disease progression. Despite these findings, the impact of interferon signaling on DN cell maturation and transitional dynamics remains unclear. To investigate the influence of IFN-α on these developmental processes, trajectory inference analysis was performed. Using the Slingshot package, the direction of B cell maturation was reconstructed across subpopulations, revealing pathway activity and metabolic reprogramming that underpin IFN-α–associated cellular dynamics.

Slingshot analysis identified five potential developmental trajectories among B cell subpopulations across all samples. Trajectory 1 connected transitional B cells to naïve IFN and naïve populations, progressing toward DN subsets via IgM memory cells **(Figure 4B**). Three additional trajectories primarily captured differentiation from naïve to classical memory compartments, including classical memory, stressed classical memory (CD69^⁺^), and memory-platelet populations. In contrast, Trajectory 5 (**Figure S6**) was specific to the DN compartment, terminating at DN2 and DN2_CXCR3^⁺^ cells, and revealing a branching pattern in which DN1 and DN4 subsets appeared developmentally closer to DN2_CXCR3^⁺^ than to DN2. To investigate the transcriptional dynamics underlying the previously observed expansion of DN cells, only Trajectories 1 and 5, those encompassing DN subsets, were selected for further analysis. Using the tradeSeq package, key genes significantly modulated along these developmental trajectories were identified. Functional enrichment analysis with EnrichR revealed that genes associated with Trajectory 1 were significantly enriched for type I interferon signaling (GO:0051607, GO:0035455), B cell activation (GO:0050864), and differentiation processes (GO:0045577). In contrast, genes along Trajectory 5 were not enriched for any specific Gene Ontology category. A heatmap of Trajectory 1 was generated to visualize the dynamics of B cell activation, IFN-α response, and stress-related pathways. Genes were selected using the associationTest() function and ordered by pseudotime across B cell subsets (**Figure 4C**). Similarly, a second heatmap was constructed to examine metabolic changes, focusing on genes involved in glucose, amino acid, and lipid metabolism (**Figure 4C**). Together, these heatmaps revealed distinct IFN-α–associated transcriptional and metabolic signatures linked to B cell activation and DN lineage expansion, supporting a role for type I interferon signaling in shaping the developmental trajectory of DN B cells.

### 3.6 B Cell–Specific IFN-**α** Scores Strongly Correlate with Ro60 and SSB Autoantibodies and Reveal Subset-Specific Links to SjD Activity

The bulk RNA-seq and flow cytometry data presented earlier, and later the single-cell RNA-seq analysis, highlight specific biological processes and B cell subpopulations that distinguish SjD patients based on B cell IFN-α stratification, findings relevant clues to explain SjD heterogeneity. To consolidate the potential role of the B cell IFN-α signature in SjD stratification, its association with autoantibodies was investigated using clinical data from the PRECISESADS cohort **(Figure 5A**). Interestingly, the B cell–specific IFN-α score showed stronger correlations with Ro60, SSB, and ENA autoantibody levels measured by ELISA, compared to the whole-blood IFN-α score derived using the Kirou method (**Figure S7**) [25]. To further investigate which B cell subtypes predominantly contribute to the B cell IFN-α signature and its relationship with autoantibody production, metadata from the single cell dataset from Arvidsson et al. 2024 were analyzed **(Figure 5B**). Mean IFN-α gene expression was calculated for each B cell subset and correlated with autoantibodies levels. A significant positive correlation was observed across all DN and the other B cell subpopulations with both SSB and Ro60 autoantibodies, except for naïve IFN B cells in the case of SSB, and for naïve IFN B cells and plasmablasts in the case of Ro52. Correlations with SSB and Ro60 were consistently positive (R = 0.48–0.65), with DN2_CXCR4, Naive and stress classical memory B cells displaying the high correlation coefficients. These same subsets also exhibited the strongest and statistically most significant associations with the Focus score in SjD patients, further supporting their potential involvement in disease activity. **(Figure 5C**)

**Figure 5:**
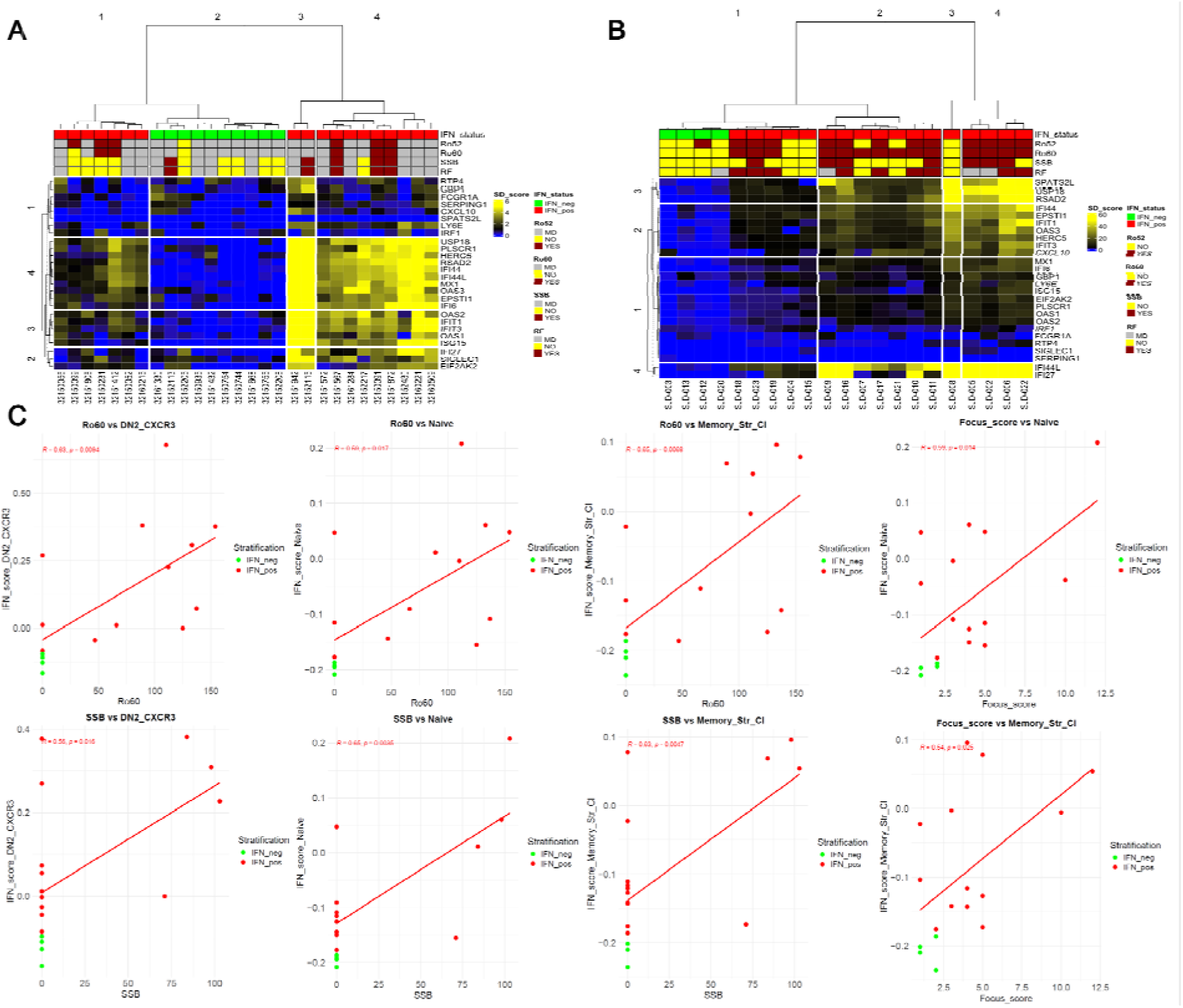
B Cell–Specific IFN-α Signatures Define and Correlate with Autoantibody Production and Focus Score. **A)** Hierarchical clustering of SjD patients from the PRECISESADS dataset based on B cell– specific IFN-α signature gene expression, as shown in Figure 2A. The heatmap displays IFN status (red: IFN-positive; green: IFN-negative) along with the corresponding autoantibody profiles for Ro52, Ro60, SSB, and RF. Columns represent individual patient samples; rows represent IFN-α signature genes. **B)** Hierarchical clustering of public single-cell RNA-seq data as shown in Figure 3A, using the B cell IFN-α gene signature. The heatmap indicates IFN status (red: IFN-positive; green: IFN-negative) and the associated autoantibody profiles (Ro52, Ro60, SSB, and RF). **C)** Pearson correlation analysis between B cell subset–specific IFN-α expression (DN2_CXCR4, naïve, and stress-responsive classical memory B cells) and Ro60, SSB autoantibody levels, as well as Focus score. IFN-neg patients are shown in green and IFN-pos patients in red. Reported on the upper-left corners of each plot are the Pearson R values and associated p-values.

## 4. Discussion

IFN-α signaling in B cell promotes their maturation and differentiation into effector B cells, thereby initiating and sustaining immune responses in autoimmune conditions [39]. In SjD, both IFN-α signaling and B cell involvement are well-established and currently utilized for patient stratification. However, a detailed characterization of IFN signaling specifically within B cells in SjD is still lacking. The present study, making use of bulk and single-cell transcriptomic analyses uncovered distinct subsets of SjD patients defined by the presence or absence of an IFN-α signature, prompting deeper investigation into the quantitative and transcriptional alterations within their B cell compartments

Interestingly, metabolic reprogramming emerged as one of the most prominent features distinguishing B cell subsets by IFN-α stratification, as well as by SjD status. Both IFN-pos and IFN-neg SjD patient B cells exhibited a shared reduction in oxidative phosphorylation (OXPHOS) activity compared to controls, with this decline observed across transitional, naïve, memory, and double-negative subpopulations. The downregulation of OXPHOS may play a critical role in modulating B cell activation via reactive oxygen species (ROS) and NFAT signaling, mechanisms well-documented in T cells, by limiting electron transport chain activity and thereby enhancing ROS production to promote immune activation [40]. These findings suggest that alterations in OXPHOS represent a pre-metabolic state independent of the IFN-α signature. This state appears to precede a more pronounced shift toward glycolytic metabolism, a requirement for plasmablast differentiation [41], which was notably evident only in the IFN-pos group. The glycolytic switch in IFN-positive B cells was broadly distributed across double-negative, classical memory, IgM memory, transitional, and naïve subpopulations. This shift was frequently accompanied by increased utilization of amino acids and lipid oxidation, indicating a comprehensive and energetically demanding metabolic reprogramming characteristic of IFN-pos B cells.

Another consistent feature observed in SjD B cells, regardless of IFN-α status, is the downregulation of translational and ribosomal processes across all subpopulations, with the exception of transitional cells. In the context of SjD, this suggests a pre-metabolic condition, similar to the reduction in OXPHOS, that appears to be independent of IFN-α signaling. The decrease in global translation may reflect cellular stress, as previously described in autoimmune settings [42,43]. In the IFN-neg group, this stress appears to primarily affect translation and, in particular, immunoglobulin production, without evidence of full activation of stress response pathways. This may indicate a condition in which stress is present but effectively managed or compensated. In contrast, IFN-pos B cells show clear signs of unresolved cellular stress, supported by the activation of apoptotic and NF-κB signaling, particularly in classical memory, DN1_ID3, and DN4 subsets, alongside upregulation of autophagy-related pathways across all subpopulations. Taken together, the coordinated reduction in both OXPHOS and translation suggests a potential pre-activated or stressed metabolic state in B cells, which may precede or accompany the active form of SjD. These alterations could represent hallmark features of the disease, independent of IFN-α status, and may offer promising avenues for clinical stratification and intervention. Independent of the pre-metabolic state, IFN-α signaling plays a fundamental role in disease progression by profoundly influencing gene expression as well as the maturation and expansion of B cell populations. Bulk and single-cell transcriptomic analyses demonstrated IFN-pos–specific upregulation of pathways related to B cell activation and proliferation, findings corroborated by flow cytometry data showing an increased frequency of activated B cells. These features were absent in IFN-neg patients when compared to controls. The expansion of the IFN-pos naïve B cell cluster identified by single-cell analysis further substantiates this IFN-restricted profile. Among B cell subsets, naïve B cells exhibited the most pronounced increase in transcript levels associated with IFN signaling, highlighting their central role in driving disease progression and autoimmune responses. IFN signaling, especially when coupled with Toll-like receptor (TLR) activation, has been shown to promote naïve B cell expansion and lower the activation threshold for autoreactive clones [44], facilitating autoantibody production as demonstrated in murine models [15]. This mechanism is consistent with the observed positive correlation between the IFN signature in naïve B cells and serum levels of Ro60 and SSB autoantibodies [38]. Moreover, the focus score correlates with the IFN-α signature within the naïve B cell compartment, suggesting that despite the predominance of plasma cells and memory CD27^⁺^ B cells in the SjD salivary gland infiltrates, IFN-driven activation of naïve B cells is required to support their generation and subsequent migration to the target tissue.

Alongside naïve B cells, DN B cells also exhibited an IFN-pos–restricted expansion in SjD patients, which was not observed in the IFN-neg group. This DN expansion was particularly notable within the DN2, DN2_CXCR3, and DN4 subsets. Trajectory analysis further demonstrated the dependence of this expansion on IFN-α signaling, confirming its direct role in driving these population changes. As previously described, DN subsets in IFN-pos patients showed distinct metabolic profiles, including increased glycolysis, lipid oxidation, and glycosylation processes, suggesting altered interactions with the microenvironment. DN2 subset was initially characterized in SLEC[45], revealing its TLR and IFN induced expansion and subsequent contribution to extrafollicular antibody production and tissue infiltration [46,47]. Although initially described in SLE, in SjD, similar IFN-driven expansion of DN B cells, including FcRL4^⁺^DN2-like populations, has been identified and may contribute to disease pathogenesis and lymphomagenesisC[20]. Existing literature has linked FcRL4^⁺^ B cells to this compartment through coexpression of CD11c, T-bet, and CXCR3. These cells have been implicated in promoting lymphoepithelial lesion formation and lymphoma development in the salivary gland [48]. In the present data, FcRL4 gene expression characterized the DN2_CXCR3 subset, and the associated high CXCR3 expression within this cluster supports the presence of FcRL4^⁺^ B cells, potentially reflecting their expansion and migration. This finding highlights a potential therapeutic target and a clinical biomarker for lymphoma risk stratification in SjD. The novel B cell–specific IFN-α score demonstrated superior performance in correlating with certain autoantibody levels compared to whole-blood IFN signatures. This likely reflects the direct link between B cell activation and autoantibody production. Although B cell isolation is not part of routine clinical procedures, limiting the immediate diagnostic applicability of the IFN-related B cell score, this metric nonetheless highlights its potential in predicting autoantibody production. This is due to its direct assessment of the cells primarily responsible for autoantibody generation. Our analysis demonstrated that high autoantibody titers consistently associate with a heightened B cell IFN response, underscoring the relevance of IFN–B cell interactions in SjD disease progression. Given the availability of novel therapies targeting B cells or type I interferons, disrupting this interaction is a promising therapeutic avenue. Moreover, our findings suggest that targeting B cell–specific signaling pathways or receptors involved in the IFN response may offer an even more strategic approach, intervening at the root of this pathological axis.

This study has several limitations, including incomplete clinical data for some patients, limited number of patients and a reliance on computational analyses of publicly available datasets. Nonetheless, in the absence of prior literature, this work represents the first effort to stratify SjD patients based on B cell IFN signatures, emphasizing their critical role in autoantibody production and disease pathogenesis. To conclude, this approach integrates interferon and B cell signatures currently utilized in both research and clinical settings and elucidates distinct interferon-related profiles in B cell subpopulations of SjD patients, revealing metabolic alterations with potential implications for disease progression. Furthermore, these findings identify novel clinical targets and biomarkers for disease monitoring and therapeutic intervention, supporting the use of current B cell–depleting treatments (Belimumab, Rituximab, Ianalumab), while also highlighting agents that directly target IFN signaling (e.g., Anifrolumab) as promising candidates.

