## Supplementary Tables and Figures for "Interferon-α–Driven Stratification of B Cell Reveals Metabolic Reprogramming of Double Negative, Naive and Transitional cell subsets and Refines Molecular Classification in Sjögren’s Disease"

| Sample ID | DN1ID3 | DN2 | DN2  CXCR3 | DN4 | Memory_Classical | Memory  IgM  ALOX5 | Memory  IgM  CD1C | Memory  platelets | Memory_Str_Cl | Memory  Str_Cl  CD69 | Naive | Naive_IFN | Plasmacell | Transitional |
| --- | --- | --- | --- | --- | --- | --- | --- | --- | --- | --- | --- | --- | --- | --- |
| CTRL_001 | 68 | 38 | 200 | 61 | 19 | 581 | 203 | 1 | 22 | 12 | 935 | 185 | 4 | 123 |
| CTRL_002 | 617 | 74 | 16 | 143 | 182 | 66 | 45 | 10 | 1817 | 28 | 5284 | 202 | 2 | 321 |
| CTRL_003 | 527 | 217 | 47 | 54 | 213 | 28 | 100 | 26 | 2085 | 19 | 5221 | 82 | 1 | 244 |
| CTRL_004 | 339 | 192 | 24 | 1075 | 436 | 1564 | 248 | 250 | 16 | 71 | 3851 | 125 | 36 | 240 |
| SJD_001 | 245 | 176 | 79 | 76 | 11 | 24 | 120 | 15 | 7 | 33 | 66 | 5075 | 0 | 55 |
| SJD_002 | 33 | 130 | 9 | 24 | 3158 | 11 | 15 | 7 | 36 | 18 | 852 | 3178 | 0 | 518 |
| SJD_003 | 105 | 166 | 2 | 1 | 1331 | 17 | 5 | 8 | 38 | 0 | 549 | 0 | 19 | 120 |
| SJD_004 | 536 | 1733 | 153 | 128 | 242 | 115 | 63 | 31 | 100 | 11 | 3860 | 67 | 2 | 563 |
| SJD_005 | 212 | 209 | 62 | 68 | 194 | 102 | 29 | 106 | 45 | 18 | 5277 | 607 | 1 | 843 |
| SJD_006 | 194 | 534 | 203 | 130 | 91 | 231 | 45 | 29 | 19 | 25 | 1490 | 6354 | 0 | 826 |
| SJD_007 | 605 | 358 | 90 | 547 | 108 | 683 | 191 | 3 | 211 | 10 | 6109 | 776 | 0 | 520 |
| SJD_008 | 1764 | 1228 | 959 | 5775 | 69 | 2300 | 489 | 30 | 570 | 32 | 1446 | 975 | 4 | 327 |
| SJD_009 | 64 | 310 | 11 | 5 | 141 | 1 | 15 | 5 | 770 | 4 | 5077 | 59 | 2 | 487 |
| SJD_010 | 121 | 91 | 23 | 108 | 1202 | 195 | 42 | 242 | 173 | 18 | 7569 | 1088 | 3 | 925 |
| SJD_011 | 322 | 66 | 79 | 125 | 258 | 256 | 61 | 46 | 52 | 11 | 5162 | 803 | 1 | 538 |
| SJD_012 | 155 | 235 | 48 | 20 | 2211 | 44 | 25 | 57 | 14 | 35 | 6977 | 31 | 1 | 246 |
| SJD_013 | 211 | 86 | 60 | 207 | 99 | 1592 | 251 | 35 | 110 | 32 | 4174 | 68 | 3 | 143 |
| SJD_014 | 15 | 41 | 18 | 4 | 94 | 21 | 53 | 24 | 3 | 8454 | 119 | 38 | 1 | 11 |
| SJD_015 | 426 | 687 | 749 | 818 | 85 | 2865 | 510 | 63 | 528 | 90 | 1799 | 73 | 0 | 301 |
| SJD_016 | 726 | 1601 | 195 | 1064 | 98 | 290 | 59 | 120 | 117 | 40 | 4440 | 1074 | 1 | 104 |
| SJD_017 | 500 | 180 | 86 | 510 | 133 | 861 | 189 | 36 | 387 | 32 | 2629 | 203 | 0 | 122 |
| SJD_018 | 272 | 184 | 105 | 208 | 144 | 553 | 156 | 26 | 276 | 17 | 7789 | 453 | 0 | 1051 |
| SJD_019 | 505 | 695 | 839 | 1357 | 74 | 1720 | 459 | 19 | 947 | 22 | 7692 | 304 | 0 | 468 |
| SJD_020 | 410 | 254 | 39 | 327 | 288 | 918 | 101 | 19 | 347 | 20 | 4745 | 43 | 5 | 255 |
| SJD_021 | 469 | 542 | 236 | 340 | 116 | 466 | 76 | 10 | 113 | 7 | 3285 | 171 | 1 | 317 |
| SJD_022 | 174 | 728 | 573 | 320 | 94 | 691 | 421 | 32 | 156 | 27 | 1512 | 2354 | 2 | 428 |
| SJD_023 | 213 | 133 | 60 | 229 | 103 | 555 | 239 | 16 | 195 | 5 | 3256 | 263 | 1 | 309 |
| SJD_024 | 0 | 0 | 0 | 0 | 1 | 0 | 0 | 1 | 1 | 0 | 7 | 0 | 25 | 2 |

**Table S1:** Number of cells per sample in single-cell data

**Table S2.** B-cell samples from SjD and controls of the PRECISESADS cohorts were excluded from further analysis based on the criteria used to filter them.

| **Individuals** | **Unfiltered samples** | **B cells with digital purity <90 % in MCP** | **Total samples analyzed** |
| --- | --- | --- | --- |
| **CTRL** | 27 samples | 4 samples | 23 samples |
| **Total SjD** | 41 samples | 13 samples | 28 samples |

**CTRL: controls; MCP: Microenvironment Cell Populations; SjD: Sjögren's disease**

| B cell subset | IFN_pos | IFN_neg | CTRL |
| --- | --- | --- | --- |
| DN1_ID3 (%) | 4.556513 | 3.22687 | 5.424974 |
| DN2 (%) | 6.007879 | 2.714087 | 1.822315 |
| DN2_CXCR3 (%) | 2.829942 | 0.545748 | 1.003847 |
| DN4 (%) | 7.506497 | 2.032818 | 4.662469 |
| Memory_Classical (%) | 4.029091 | 14.39089 | 2.973068 |
| Memory_IgM_ALOX5 (%) | 7.595252 | 9.416893 | 7.83141 |
| Memory_IgM_CD1C (%) | 1.953247 | 1.399165 | 2.084645 |
| Memory_platelets (%) | 0.524229 | 0.435866 | 1.003847 |
| Memory_Str_Cl (%) | 2.997874 | 1.864332 | 13.78104 |
| Memory_Str_Cl_CD69 (%) | 0.247109 | 0.318658 | 0.454704 |
| Naive (%) | 44.21401 | 60.23368 | 53.48374 |
| Naive_IFN (%) | 12.00554 | 0.520108 | 2.07765 |
| Plasmacell (%) | 0.011493 | 0.102557 | 0.150402 |
| Transitional (%) | 5.521324 | 2.79833 | 3.24589 |

**Table S3:** Percentage of B cell subset within IFN-pos, IFN-neg, and CTRLs groups.

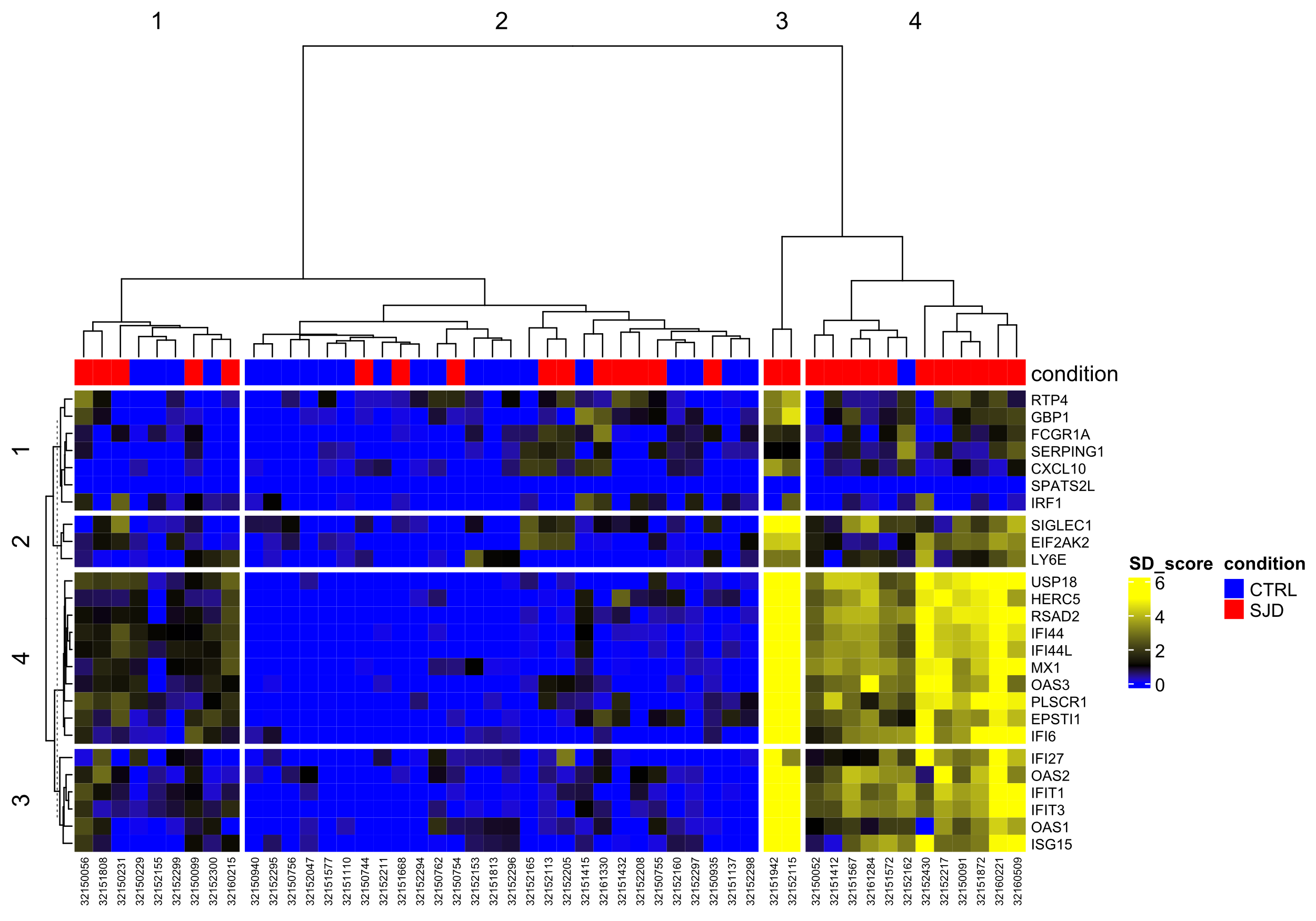

**Figure S1. A)** Hierarchical clustering of SjD patients and CTRLs based on the B cell IFN-α signature from the PRECISESADS dataset. The heatmap displays conditions: SjD patients are shown in red, and CTRL in blue. Columns represent individual samples; rows correspond to IFN-α signature genes. Each cell indicates the gene expression Z-score relative to the CTRL group.

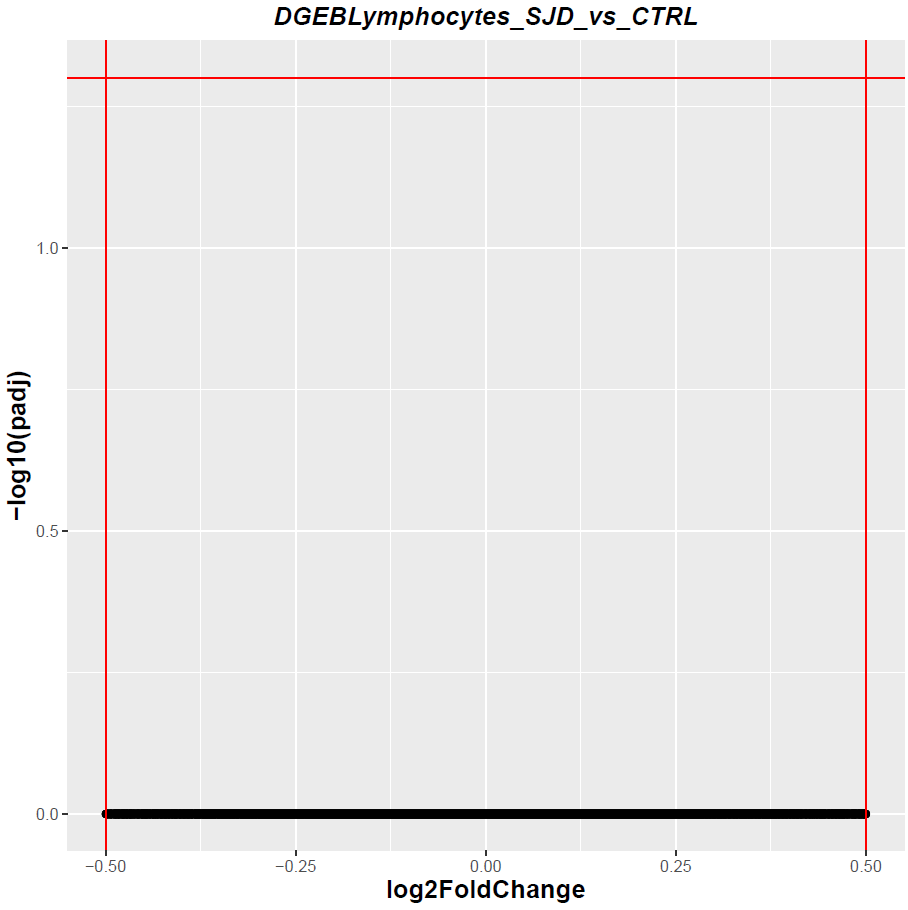

**Figure S2**. Volcano plot of the differential gene expression analysis comparing IFN-neg SjD patients to CTRLs in the PRECISESADS bulk B cell dataset. Significantly upregulated genes are shown in red, and downregulated genes in blue. The top 25 genes in each direction are labeled. Red lines indicate significance thresholds: |log₂FC| > 0.5 and false discovery rate (FDR) < 0.05

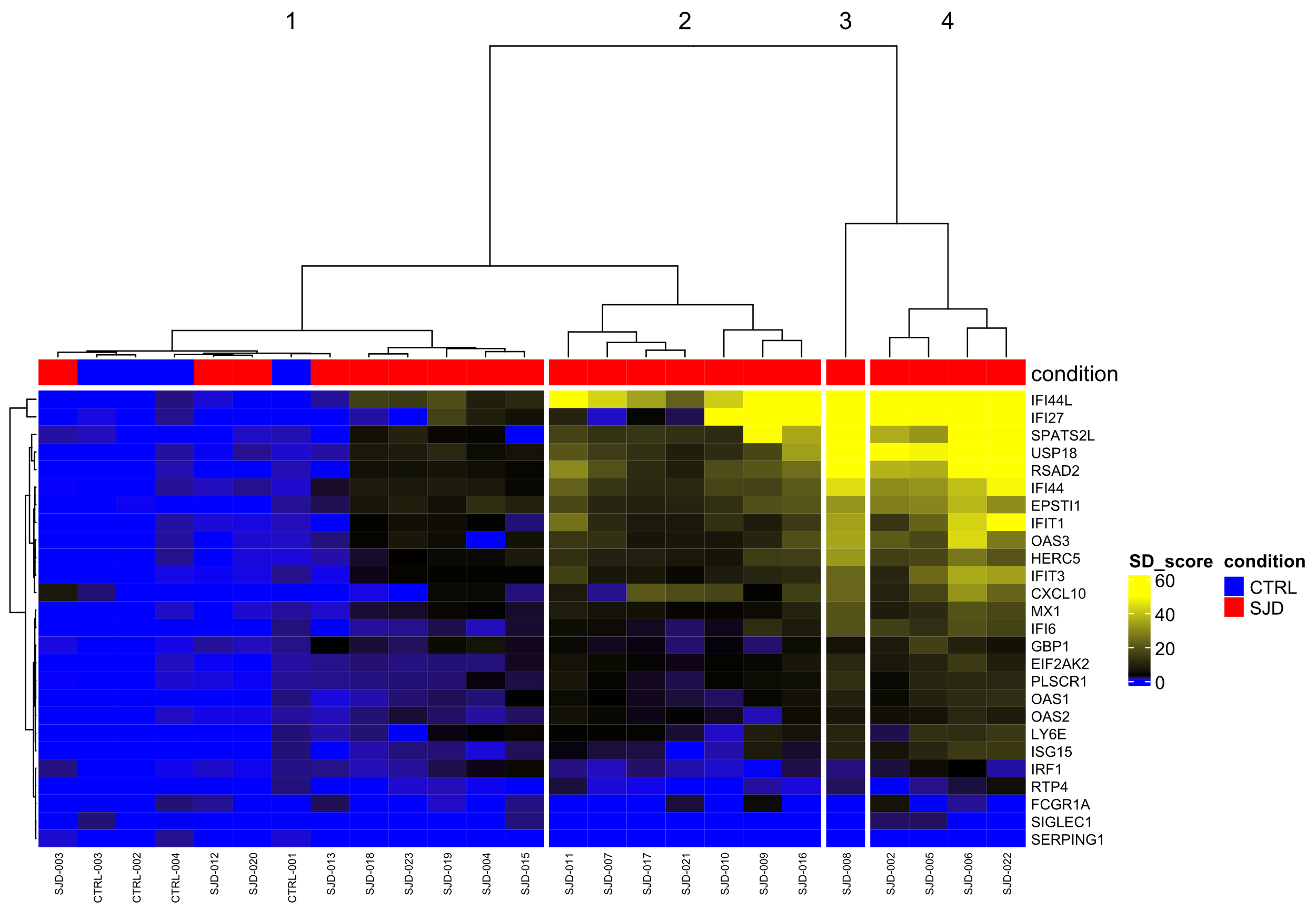

**Figure S3. A)** Clustering of public single-cell data by hierarchical clustering. The heatmap displays conditions, with SjD patients in red and CTRLs in blue. Columns represent individual samples; rows represent IFN-α signature genes. Each cell shows the gene expression Z-score relative to the CTRL group.

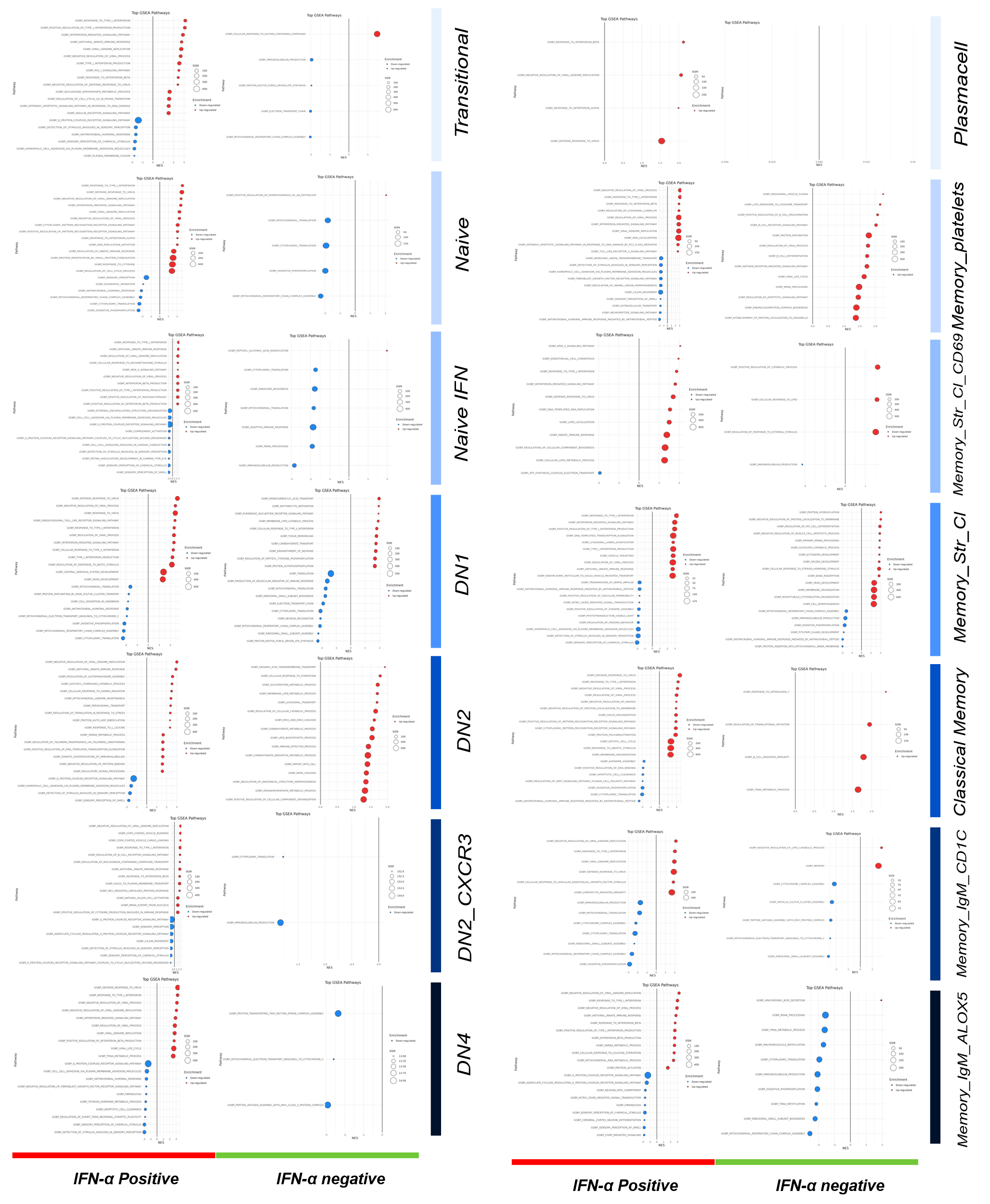

**Figure S4**: GSEA analysis on the B cell subsets among IFN-α groups and CTRLs. The GSEA pathways that increased and decreased are, respectively, shown in red on the right and in blue on the left. On the Y axis, the GSEA results are divided by B cell subpopulations, including Transitional, Naive, Double negative, Classical memory, and IgD+ Memory B cells. On the X axis, the GSEA results are stratified by the groups: IFN-positive, IFN-negative, and CTRLs.

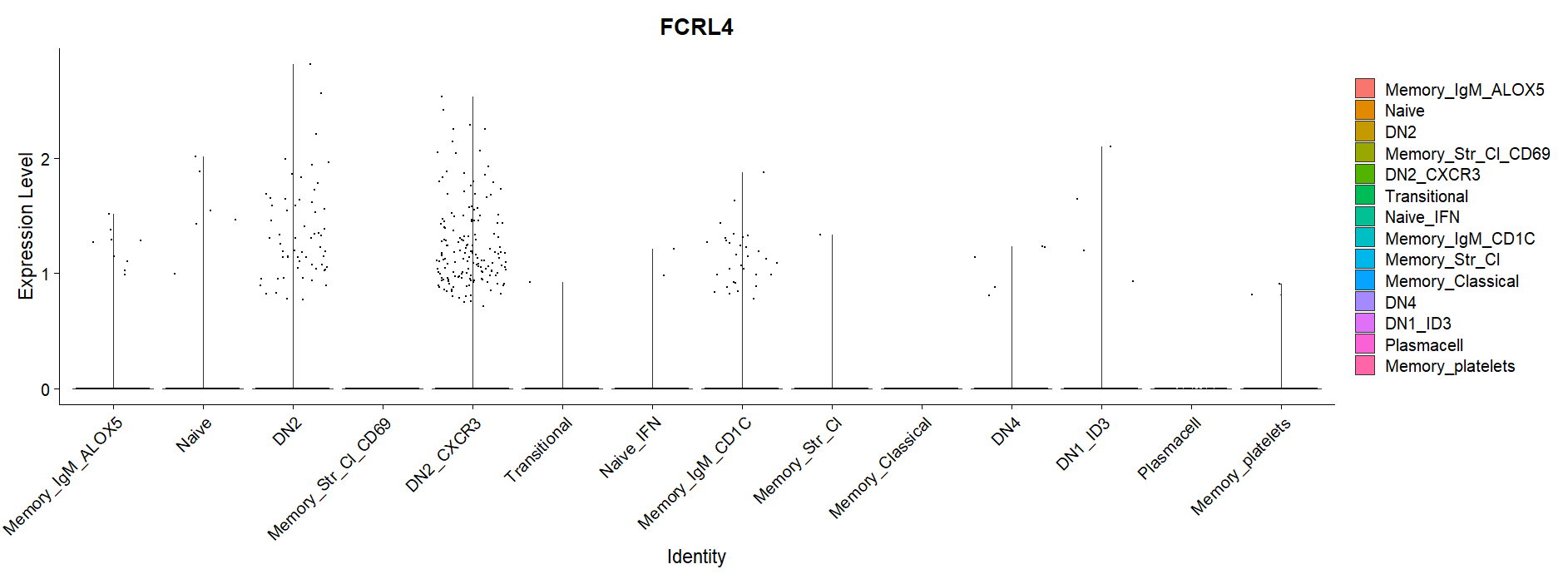
**Figure S5**: Violin plot showing the expression of FcRL4 in the B cell subpopulation of the public single cell data. The violin plot confirms the specific expression of the DN2_CXCR3 subpopulation.

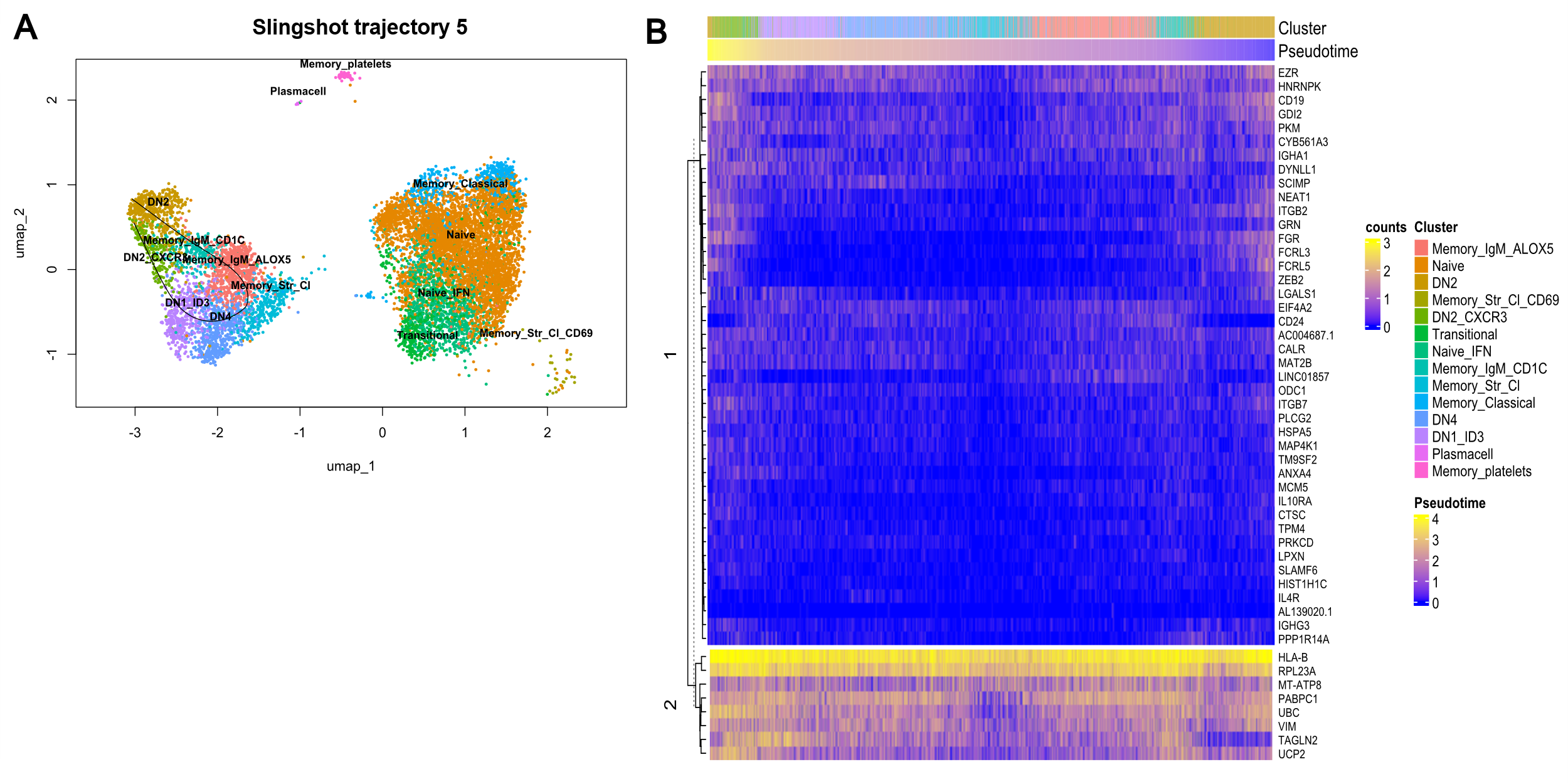

**Figure S6 A)** Trajectory analysis of B cell maturation using Slingshot on Lineage 5, visualized on a UMAP plot with inferred developmental progression. **B)** Heatmap of B cell subtypes ordered by pseudotime along Trajectory 1. The x-axis shows the 50 top-most significant genes identified as trajectory-relevant by associationTest().

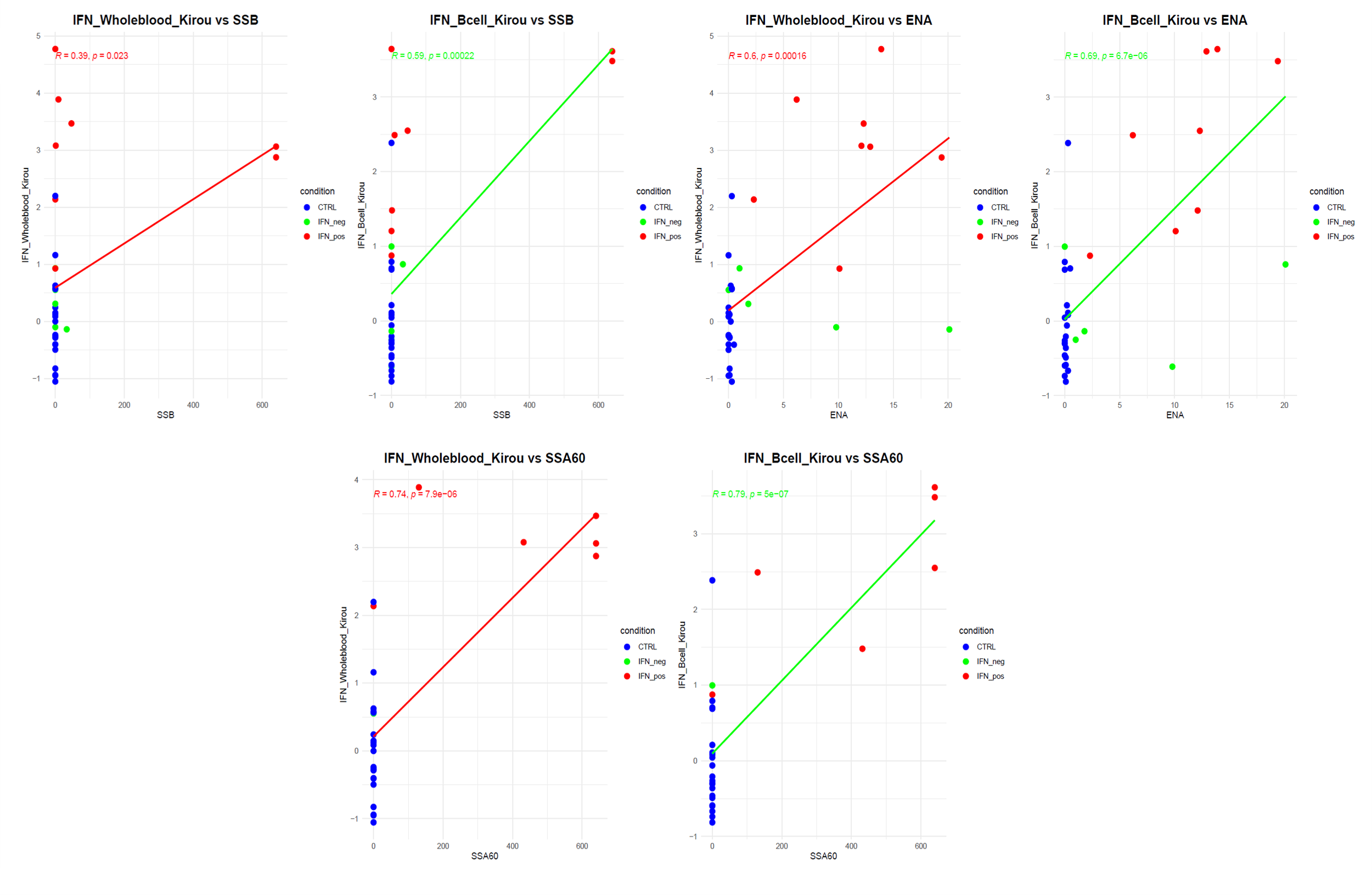

**Figure S7 A)** Correlation between Ro60, SSB, and ENA autoantibody levels (X-axis, measured by ELISA) and IFN-α scores: whole-blood IFN-α score calculated using the Kirou method (red line) and B–cell–specific IFN-α score (green line) shown on the Y-axis.
